# Simulating the diffusion of PPR-like disease to improve surveillance

**DOI:** 10.1101/2024.12.08.627410

**Authors:** Asma Mesdour, Sandra Ijoma, Muhammad-Bashir Bolajoko, Mamadou Ciss, Eric Cardinale, Stephen Eubank, Mathieu Andraud, Andrea Apolloni

**Author notes:** these authors contributed equally to this work (Mathieu Andraud, Andrea Apolloni).

## Abstract

Animal mobility is a founding element of pastoral culture and fundamental to the economy in West Africa. Herds’ movement, involving mainly cattle, sheep, and goats, is vital for adapting to climate fluctuations, optimizing natural resources, and managing risks in livestock production, as well as livestock trade and exchanges. However, these movements contribute to the spread of transboundary animal diseases such as Peste des petits ruminants (PPR). Nigeria experienced its first PPR outbreak in the 1960s-1970s, and outbreaks have regularly occurred since then. Yet, no adequate surveillance system has been put in place, and the absence of proper animal movement tracking remains a critical shortcoming in addressing this ongoing threat. Because of this, we rely on *ad-hoc* activities, like mobility surveys, to collect this information. However, these data could be partial and limited in time and space, hindering the capacity to identify suitable areas for monitoring disease circulation (*sentinel nodes*). Market survey data from three northern states are collected once to reconstruct the small ruminant mobility network. A group of areas with a high potential to infect each other were identified (Contagion Cluster). A stochastic Susceptible-Infected-Recovered (SIR) was used to simulate the spread of a PPR-like disease through movements. Sentinel nodes and their key characteristics were identified using Random Forest classification. The network (missing movement) was predicted with a hierarchical random graph (HRG), and uncertainty analysis assessed the effects of missing movements on the epidemic extent and identity and characteristics of sentinel nodes. The number of sentinel nodes varied with the epidemic’s severity, but their characteristics remained consistent. The uncertainty analysis results showed that adding 1% of the most probable missing links did not affect the final size of the epidemic. However, a significant difference appears when adding more than 3% of the missing links, causing a gradual fluctuation in the epidemic’s final size. The final size of the epidemic stabilized when adding more than 50% of the most probable links. The challenge posed by incomplete animal movement data underscores the need for further research and data collection efforts. However, predicting missing links presents a promising method for enhancing the reliability of epidemic prediction and sentinel node identification. This potential to improve disease surveillance and control strategies is a significant implication of our study.

## Introduction

Transboundary animal diseases (TADs) are highly contagious diseases that can rapidly spread worldwide, posing significant socioeconomic and public health risks. A deeper understanding of their transmission and spread is essential^1^. Among these diseases, Peste des petits ruminants (PPR). It threatens over 68% of the world’s small ruminant population^1^ and the livelihood of more than 300 million smallholders^2^. PPR is a severe viral disease that predominantly affects domestic and wild ruminants; goats and sheep are particularly susceptible^3^. In naive sheep and goat populations, morbidity and mortality can be greater than 90%^4^. In enzootic areas, morbidity and mortality vary between 10 and 100%^5^. The Peste des Petits Ruminants virus (PPRV) can be easily transmitted through direct contact with secretions and excretions from infected animals and through contact with contaminated objects, known as fomites^6^. Despite the availability of an inexpensive and effective live-attenuated vaccine, the virus continued to spread^7^.

The main factor contributing to this rapid spread is the movement of animals^8,9^. Animal movement control is therefore one of the key components for infectious disease control and eradication programs^3^. Given that the Food and Agriculture Organization (FAO) and the World Organization for Animal Health (WOAH) aim to eradicate PPR by 2030, surveillance of this disease is a priority and should be improved^2^. Animal mobility is a complex phenomenon, and monitoring the spread of disease through it requires the use of tools such as complex networks^10–12^ combined with mathematical modeling^13,14^, using classical models such as Susceptible-Infected-Recovered (SIR) enables the simulation of the spread of infectious animal diseases such as PPR.

Accurate recording of animal mobility data enables effective identification of disease transmission hotspots or the most vulnerable spots; This information strengthens targeted surveillance and control measures, improving the ability to prevent and manage outbreaks. Access to detailed data requires systematic collection of animal movement patterns, which requires considerable logistical and financial resources. To date and to our knowledge, several countries vulnerable to PPR have no system to identify these movements, such as Nigeria, where numerous outbreaks of PPR have been described since the 1960s-1970s^8^. Therefore, identifying vulnerable areas without detailed data or with only fragmented data is a complex task. Using link (animal movement between two areas) prediction models based on a fraction of surveyed samples is helpful in low-income countries, where this data is generally unavailable and the cost-benefit ratio is less convincing, and even very fragmented information can be used to deduce the network structure^15^.

Surveillance not only involves identifying vulnerable areas, but can also be extended to identifying groups of areas that share similar vulnerability to disease and with a high risk of rapid infection and spread to commercial partners. Nath et al. (2019) and Mishra et al. (2023)^16^,^17^ developed a method capable of identifying contagion clusters, which they define as connected maximal subsets of nodes such that if the infection is introduced to the cluster, it spreads rapidly within the cluster. This approach is useful in our context for optimal monitoring. Furthermore, in previous studies^18^, we hypothesized that the backbone (a simplified network that contains only the most important edges)^19^ could serve as a good candidate for surveillance, but its impact on the final size of the epidemic has not been thoroughly investigated.

As previously mentioned, Nigeria faces a significant challenge with Peste des petits ruminants (PPR), worsened by the lack of a reliable system to track animal movements. This makes it crucial to identify the most vulnerable areas for effective surveillance quickly. Although we only have a portion of the data from 10 markets in 3 northern states^20^, our study aims to predict missing connections (movements not captured due to a non-exhaustive and irregular survey) from these partial data to better understand where animals move from and to. We then simulate the spread of PPR. The results obtained - specifically the final maximum size of the epidemic, the peak of the epidemic, and the infection frequency of each area - are used to identify vulnerable zones. By comparing our results with observed and predicted data, we can assess whether our limited data is sufficiently informative to draw solid conclusions and achieve reliable outcomes, as suggested by Charters et al. (2019)^15^. Furthermore, to optimize surveillance, our aim is to identify groups with similar vulnerabilities and to verify whether previously identified vulnerable areas exist within these communities. Then, we explore the role of vulnerable areas and routes (called “bridges” in networks) connecting different clusters, and backbone by assessing their impact on the final epidemic size.

The present study presents a modeling work to better understand the spread of the PPR virus in 3 northern states of Nigeria and to assess the impact of control measures. In the first step, we propose to identify potential contagion clusters from movement data. Then, a PPR simulation model was developed, based on a SIR approach to the observed network, to identify vulnerable areas with their structural and/or socio-economic characteristics. Surveillance and control strategies were then tested through the removal of nodes and edges in vulnerable areas and backbone and the removal of inter-cluster links. Finally, an uncertainty analysis was performed to assess the robustness of the results using a predicted network based on the partially observed one.

## Results

### Communities and PPR simulation on Nigeria’s animal mobility network

#### Cluster contagion are geographically dispersed

The network used in this study is detailed in^20^. Briefly, we used the village level to represent our nodes (nodes = 235), with the links (movements between villages) weighted by the number of animals moved (links = 355). Figure 1 illustrates the contagion clusters for the animal mobility network. Each cluster is characterized by a size, *n*, corresponding to the desired final size of the strongly connected component (SCC) (n = 2, n = 5, 10, n = 20, n = 30, n = |V|), where |V| represents the total number of nodes in the network (|V|=235). The colored clusters represent the contagion clusters identified using the COCLEA algorithm (COntagion CLusters Extraction Algorithm)^16^,^17^. Nodes that do not belong to any of these clusters are not shown. The COCLEA algorithm revealed the presence of geographically dispersed communities, regardless of the chosen value of n. In this study, we selected n = 20 as it provides a good balance between n = 10 and n = 15 (Figure 1). At n = 15, nodes in the South are excluded but included at n = 20. The network reveals seven communities distributed across 17 states in the North and South of the country. The largest community comprises 16 nodes (distributed in 8 states: Bauchi, Edo, Federal Capital Territory, Jigawa, Kaduna, Kano, Plateau and Rivers), followed by communities with 15 (distributed in 7 states: Bauchi, Gombe, Kaduna, Kano, Nasarawa, Plateau, and Yobe), 13 (distributed in 9 states: Bauchi, Benue, Federal Capital Territory, Kaduna, Kano, Lagos, Ondo, Plateau and Yobe), 6 (distributed in 4 states: Bauchi, Federal Capital Territory, Jigawa and Kano), 3 (distributed in 2 states : Kano and Yobe), and two nodes each (Two nodes were distributed in Ebonyi and Bauchi, and two additional nodes in Katsina and Kano). Bauchi, Kano, the Federal Capital Territory (FCT), Plateau, and Yobe are the most frequent states, appearing in multiple communities. Additionally, 181 nodes do not form communities and remain isolated.

**Figure 1.**
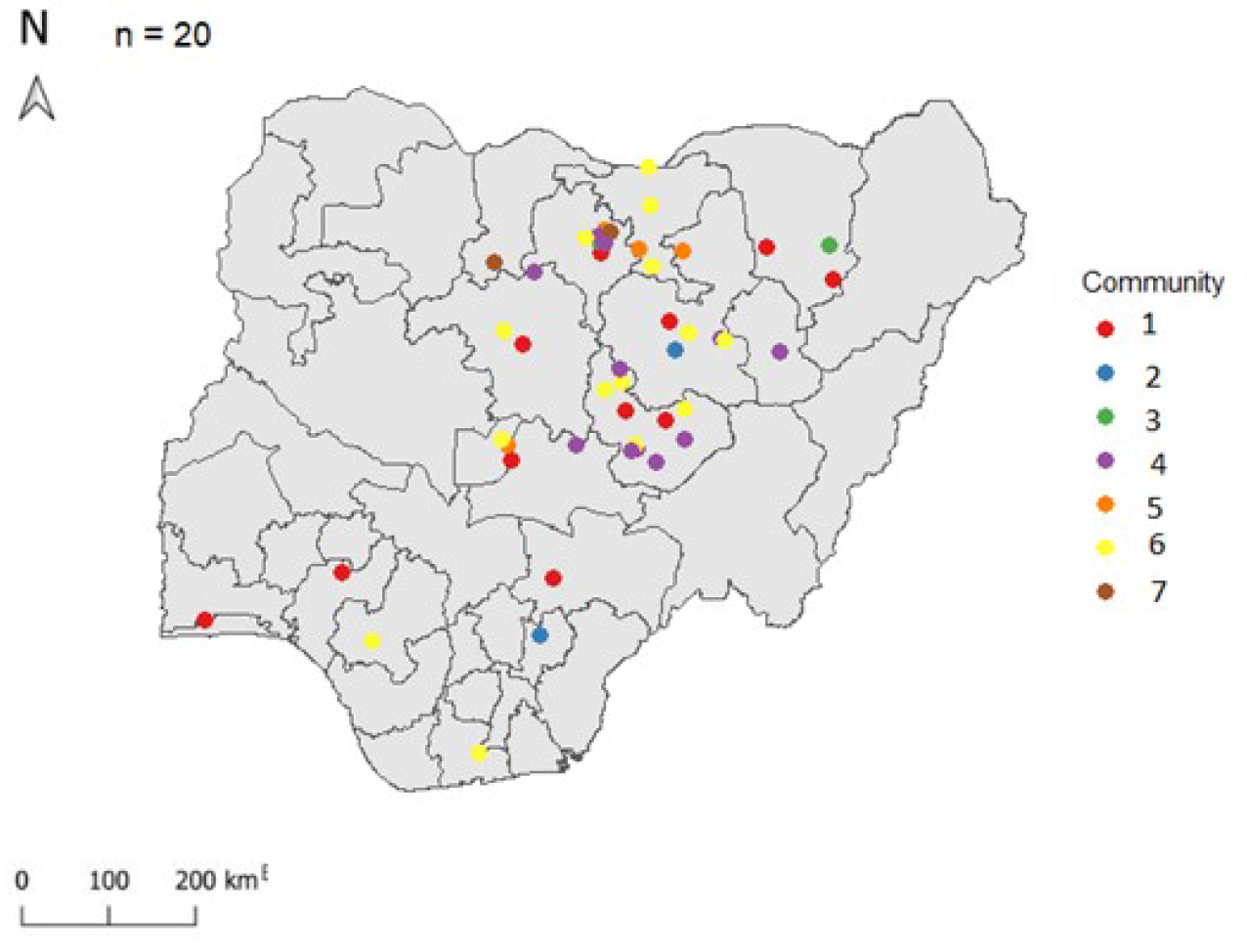
Contagion cluster identified using COCLEA algorithm: a group of a nodes with the same potential to infect each other. The map represents the contagion cluster with n = 20. Seven clusters were identified (color).

#### Sensitivity analysis of transmission (Pinf) and recovery probability (Prec) combinations

In this study, we examined 100 values of *Pinf* (between 0.00001 and 0.2) combined with 100 values of *Prec* (between 0.001 and 0.5), resulting in 10,000 unique combinations. Twelve of the 10,000 combinations did not trigger an epidemic; therefore, they were excluded from further analysis. Among the 9,988 remaining combinations, four groups (Q1, Q2, Q3, and Q4) were selected based on their average final epidemic size, as simulated using the weighted SIR model, with 300 simulations conducted for each combination (Figure 2).

**Figure 2.**
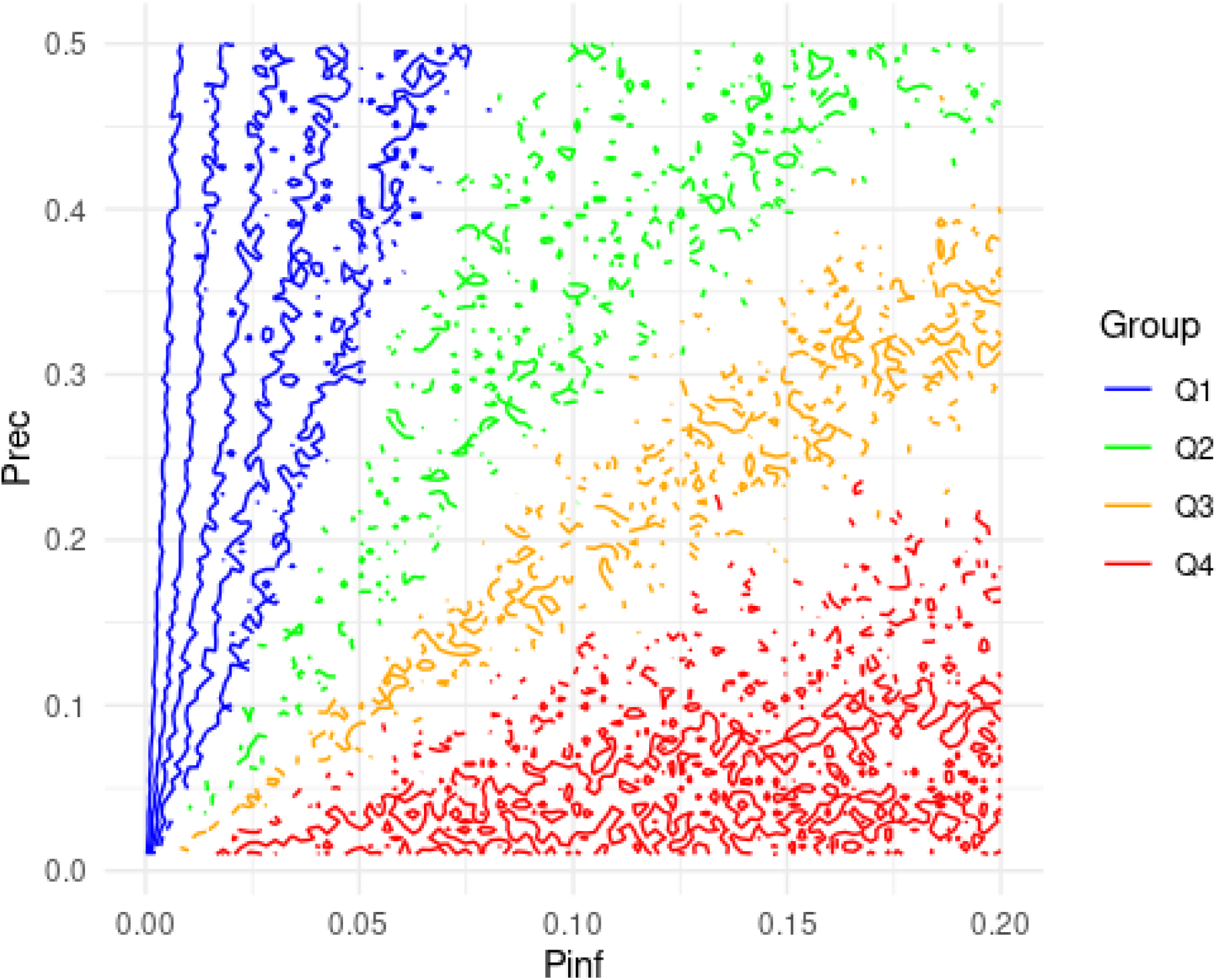
Contour plot of the sensitivity analysis: x represents *Pinf* values range from 0.00001 to 0.2, y represents *Prec* values range from 0.001 to 0.5. The color represents the performance (average final size) obtained from the SIR weighted model

Figure 3 highlights the distinct characteristics between groups Q1 to Q4, based on their average final epidemic size and the average time to reach the epidemic peak. Group Q1, characterized by *Pinf* values ranging from 0.00001 to 0.11 and *Prec* values from 0.01 to 0.5, exhibits the smallest final epidemic size, with an average of only 18 infected nodes. In contrast, Group Q4, with *Pinf* values between 0.012 and 0.2 and *Prec* values from 0.01 to 0.34, shows the largest final epidemic size, averaging 49 infected nodes. Furthermore, the epidemic peak occurs later in Group Q4, averaging 4.92 weeks compared to 3.53 weeks in Group Q1. Groups Q2, with *Pinf* values ranging from 0.004 to 0.2 and *Prec* values from 0.01 to 0.5, and Q3, with *Pinf* values ranging from 0.006 to 0.2 and *Prec* values from 0.01 to 0.5, exhibit intermediate trends in both final epidemic size—33 and 39 infected nodes, respectively—and the timing of the epidemic peak, occurring at 3.86 and 3.14 weeks, respectively. The similarity between these two groups is mainly due to the overlap in certain *Pinf* and *Prec* values.

**Figure 3.**
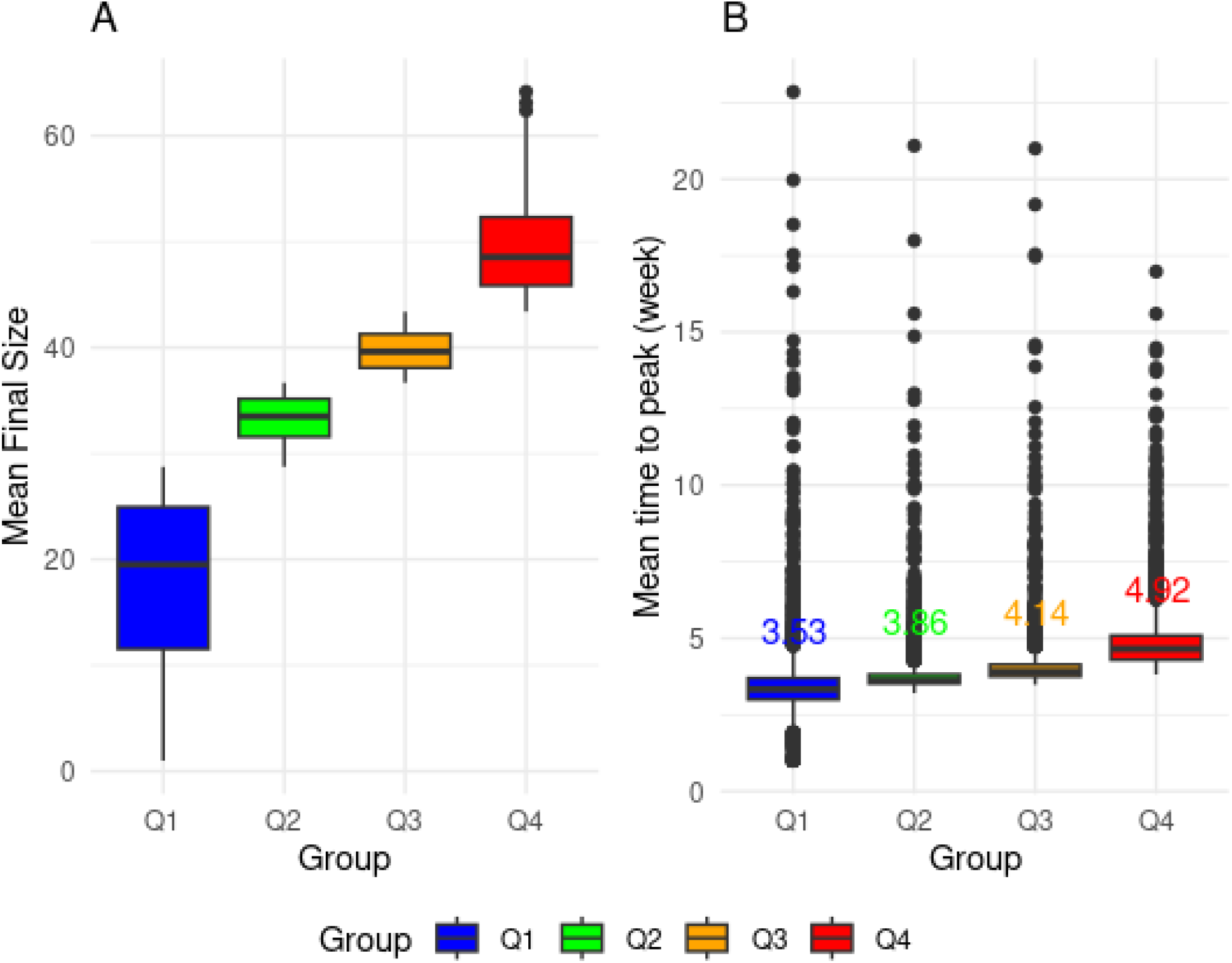
Result of PPR simulation using stochastic SIR weighted model. A: Average final size per group. Each boxplot represents the average epidemic final size of the group. Blue represents Q1 (*Pinf* = c(0.00001 : 0.11) combined to *Prec* = c(0.01 : 0.5)). Green represents Q2 (*Pinf* = c(0.004 : 0.2) combined to *Prec* = c(0.01 : 0.5)). Orange represents Q3 (*Pinf* = c(0.006 : 0.2) combined to *Prec* = c(0.01 : 0.5)). Red represents Q4 (*Pinf* = c(0.012 : 0.2) combined to *Prec* = c(0.01 : 0.34)). B: The mean time (per week) to peak.

#### Characteristic of Sentinel nodes remain stable according to the group

The number and identity of sentinel nodes were identified for each parameter group. Node statuses were determined based on how frequently each node was affected before the epidemic peak. We focus on the most vulnerable nodes—those often infected before the peak—which we call “Highly vulnerable” and consider potential sentinel nodes. Nodes affected less frequently (below a threshold set based on total frequency) are classified as “Low vulnerable” and are not considered sentinel nodes. If a node is frequently infected after the peak, it is regarded as infected but not vulnerable.

Figure 4 shows that in group 1, there are nine nodes with high vulnerability, primarily located in northern States: 2 in Bauchi, 1 in Borno, 2 in Jigawa, 1 in Nasarawa, and 3 in Plateau. Additionally, 67 nodes exhibit low vulnerability, predominantly found in Bauchi (26) and Plateau (30). In group 2, only one node with high vulnerability is identified in Kano State, while 97 nodes have low vulnerability, mostly in Bauchi (27) and Plateau (38). In group 3, 9 high-vulnerable nodes are located across Nigeria: 3 in Bauchi, 2 in Plateau, 1 in Delta, 1 in Enugu, 1 in Lagos, and 1 in Rivers. Additionally, 130 nodes display low vulnerability, mainly distributed in Bauchi (29), Kano (23), Plateau (39), and Jigawa (9). Furthermore, in group 4, an increased number of nodes become highly vulnerable (43): 1 in Abia, 1 in Anambra, 11 in Bauchi, 2 in Delta, 2 in Enugu, 2 in the Federal Capital Territory, 2 in Lagos, 5 in Nasarawa, 1 in Ondo, and 12 in Plateau. Conversely, 131 nodes exhibit lower vulnerability, similar to those in group 3, primarily located in Bauchi (30), Kano (23), Plateau (39), and Jigawa (9). As the severity of the disease increases (*Pinf ≥* 0.006), more highly vulnerable nodes emerge, predominantly located in the southern part of the country (as seen in groups Q3 and Q4). Conversely, when the disease is less severe, the first vulnerable nodes appear in the northern region (as observed in groups Q1 and Q2).

**Figure 4.**
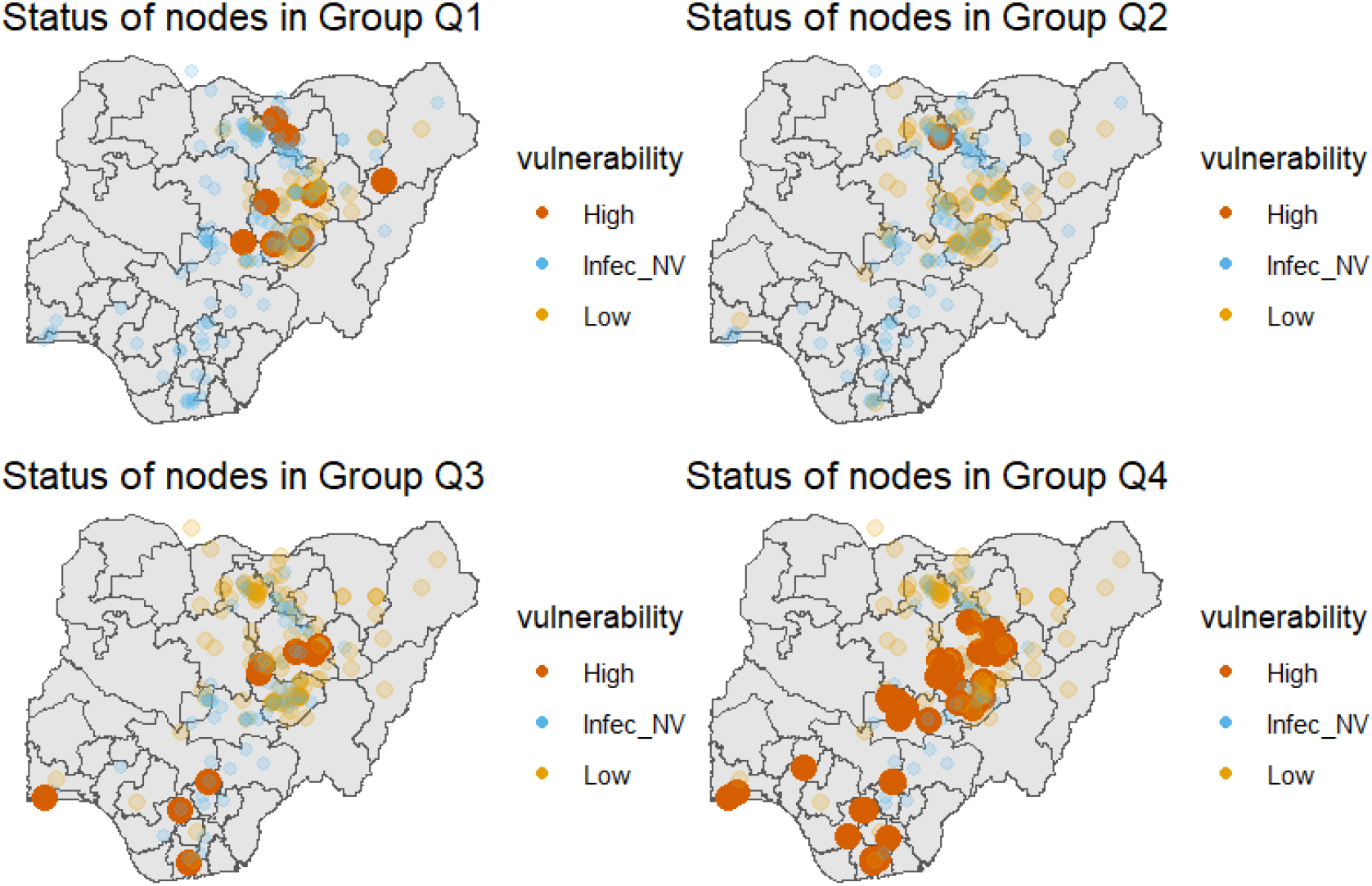
Vulnerability Distribution of Node Status (High, Infected but Not Vulnerable, Low) Across Groups Q1, Q2, Q3, and Q4 in Nigeria: Nine Highly Vulnerable Nodes in Q1, One in Q2, Nine in Q3, and Forty-Three in Q4

The identity of highly vulnerable nodes differs between group 1 and groups 2, 3 and 4. However, the identity of the highest vulnerable nodes remains consistent between groups 3 and 4 (the nine vulnerable nodes are also vulnerable in group 4). Structural characteristics are more significant in the fourth group than socio-economic and environmental factors. In-degree, incloseness, in-neighbourhood, and eigenvector remain consistent across all four groups. However, their relative importance varies, particularly between groups 1 and 2 compared to groups 3 and 4 Figure 5).

**Figure 5.**
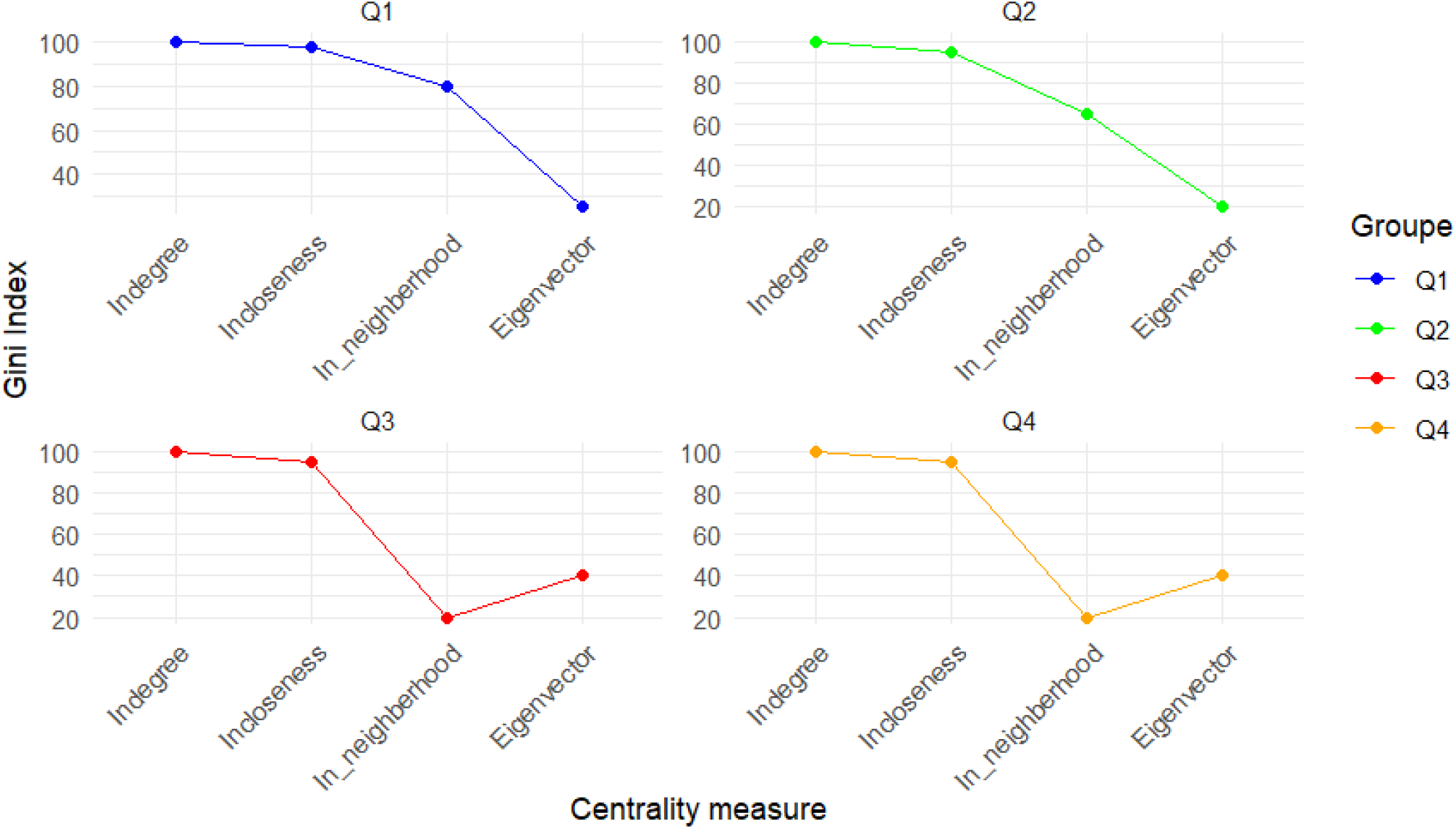
The most important characteristics of vulnerable node in each group: Result of Random Forest Classification using centrality measure (in/out degree, in/out closeness, betweenness, eigenvector, in/out strength, in/out neighbourhood, in/out H index) and socio-economic and environmental features (Human population, animal population, Dry Matter Product (DMP), Accessibility

### Impact of target surveillance: Eliminating backbone nodes drastically reduces PPR propagation

We consider a surveillance and control scenario in which once infected animals are detected at specific nodes, movement restriction measures are put in place, and animals can not move to other nodes, or animals in the market are vaccinated. In this way, the infection can not spread once detected. Scenarios are evaluated by simulating surveillance through the following actions: i) removing the set of vulnerable nodes identified in each parameter group, ii) eliminating the bridges or links that connect the previously identified communities, and iii) removing the nodes that form the network’s backbone. This backbone was extracted using disparity filter^19^.

The bottom metric offers valuable insights into how different networks contribute to its overall structure, performance, and resilience: The Largest Connected Component (LCC), often referred to as the “giant cluster,” represents the maximum number of interconnected nodes within a network^21^. Efficiency (Eff) is a measure of a network’s functional effectiveness, calculated based on the shortest paths between nodes. In weighted networks, efficiency is derived from the minimum sum of the inverse link weights, enabling the identification of stronger links as more direct routes^22^. Higher efficiency indicates that nodes are closely connected, while a decrease in efficiency signifies longer shortest paths between nodes^21^. Total Flow (TF) is the overall quantity of animals moving through the connections or pathways within the network. Table 1 illustrates that removing nodes classified as vulnerable from groups Q1 and Q4 has a minimal impact on these measures compared to the reference network. Removing vulnerable nodes from groups Q2 and Q3 has a more significant impact, especially in Q2. Although essential, bridges are not a significant part of the overall structure. The backbone is highly critical to network connectivity. Its removal drastically reduces the LCC, decreasing the number of nodes from 229 to 25 and the number of links from 333 to 28 (Table 1). The network shows resilience when vulnerable nodes from groups Q1 and Q4, as well as bridges, are removed, with minimal impact on overall efficiency. In contrast, removing nodes from group Q2 enhances efficiency, while removing nodes from group Q3 decreases it. The backbone is critical for network efficiency; its removal significantly lowers overall efficiency from 1.32*e*^*™*03^ in the reference network to 5.95*e*^*™*05^ in the network without the backbone. Removing vulnerable nodes in groups Q2 and Q3 reduced the total flow. Removing the bridges had less impact on total flow than groups Q2 and Q3. The backbone is crucial for maintaining total flow through the network; its removal drastically decreased the total flow from 29,992 in the reference network to 2,948 without the backbone.

**Table 1.** Impact of suppression of node or link on robustness of networks: Six networks are generated according to the scenarios of surveillance: network without high vulnerable node of Q1, network without high vulnerable node of Q2, network without high vulnerable node of Q3, network without high vulnerable node of Q4, network without bridge linking the communities, network without node of backbone. LCC: Largest Connected Component, Eff: Efficiency, TF: Total Flow

| Network | LCC | Eff | TF |
| --- | --- | --- | --- |
| Reference | 333 | 1.32e-03 | 29992 |
| without set Q1 | 331 | 1.33e-03 | 29903 |
| without set Q2 | 289 | 1.54e-03 | 24347 |
| without set Q3 | 318 | 8.62e-04 | 28863 |
| without set Q4 | 326 | 1.29e-03 | 29767 |
| without bridge | 327 | 1.32e-03 | 29601 |
| without backbone | 28 | 5.95e-05 | 2948 |

Figure 6 shows no notable difference in the final epidemic size when vulnerable nodes from groups Q1, Q2, and Q4 are removed compared to the reference network. Similarly, eliminating bridges does not have a major impact on the final epidemic size compared to the reference one. Eliminating vulnerable nodes from group Q3, consisting of only one node, has a negligible impact on the final epidemic size but is higher than the other groups. Removing backbone nodes has the most considerable impact on the final epidemic size, drastically reducing it. On average, one node (village) will be infected in all groups, if we prevent all disease propagation from these nodes.

**Figure 6.**
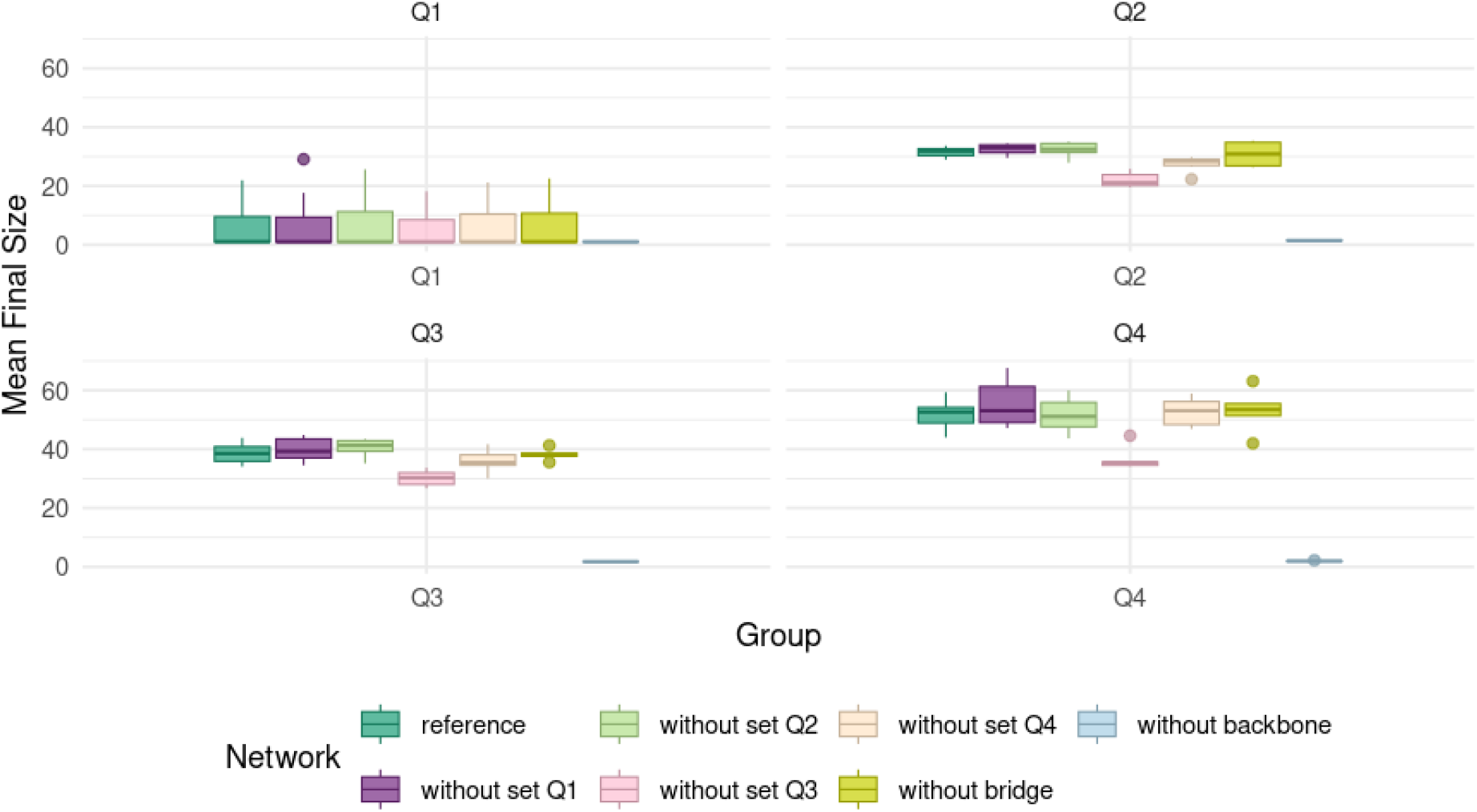
Impact of Removing Vulnerable Nodes, Bridges, or Backbone on PPR Propagation: After simulating PPR using the weighted SIR model across each network, the results were compared to the reference network regarding final epidemic size. Each plot represents a parameter group, ranging from Q1 to Q2, with different colors indicating various surveillance scenarios.

### Uncertainty study

#### variation of the final size epidemic according to the number of added links

The Hierarchical Random Graph (HRG) model is used to predict missing links (AUC = 0.9). This model searches the space of all possible dendograms of a network for the ones that best fit its hierarchical structure. Non-adjacent node pairs with a high average probability of being connected within these dendograms are considered good candidates for interaction. This model predicted 4670 additional links. Each link pair is assigned by its probability of existence. Considering the reference network (i.e., the observed) as a starting point, we added additional links based on their likelihood of existing. We consider two scenarios:

- *Top down scenario* Links are added in sequence from the most likely of existence to the less likely to exist
- *Bottom up scenario* Links are added in sequence from the less likely to the most likely (from 1% to exist

We ran PPR simulations on each of the generated networks and compared the final size of the epidemic with the reference network. To facilitate comparison of the results, we have fixed (*Pinf* at 0.1 and varied (*Prec* between 0.01 and 0.5.

In the *Top down scenario*, we observed that the average final size of the epidemic behaves similarly across the networks for each value of (*Prec* (Figure 7). Starting with the reference network, adding 1% of the most probable links is sufficient to change the average final size of the epidemic when (*Prec* = 0.01. However, as the (*Prec* (from 0.13 to 0.5) increases, the final size is not significantly different (p-value=0.238). However, with the addition of 3% of links, the final size of the epidemic becomes significantly different for different *Prec* (p-value=0.0059). For the other percentages of added links (5%,8%, 10%, 20%, 30%, 40%, 50%, 60%, 70%, 80%, 90%, 100%) the respective p-values are shown in Table tab:table

**Figure 7.**
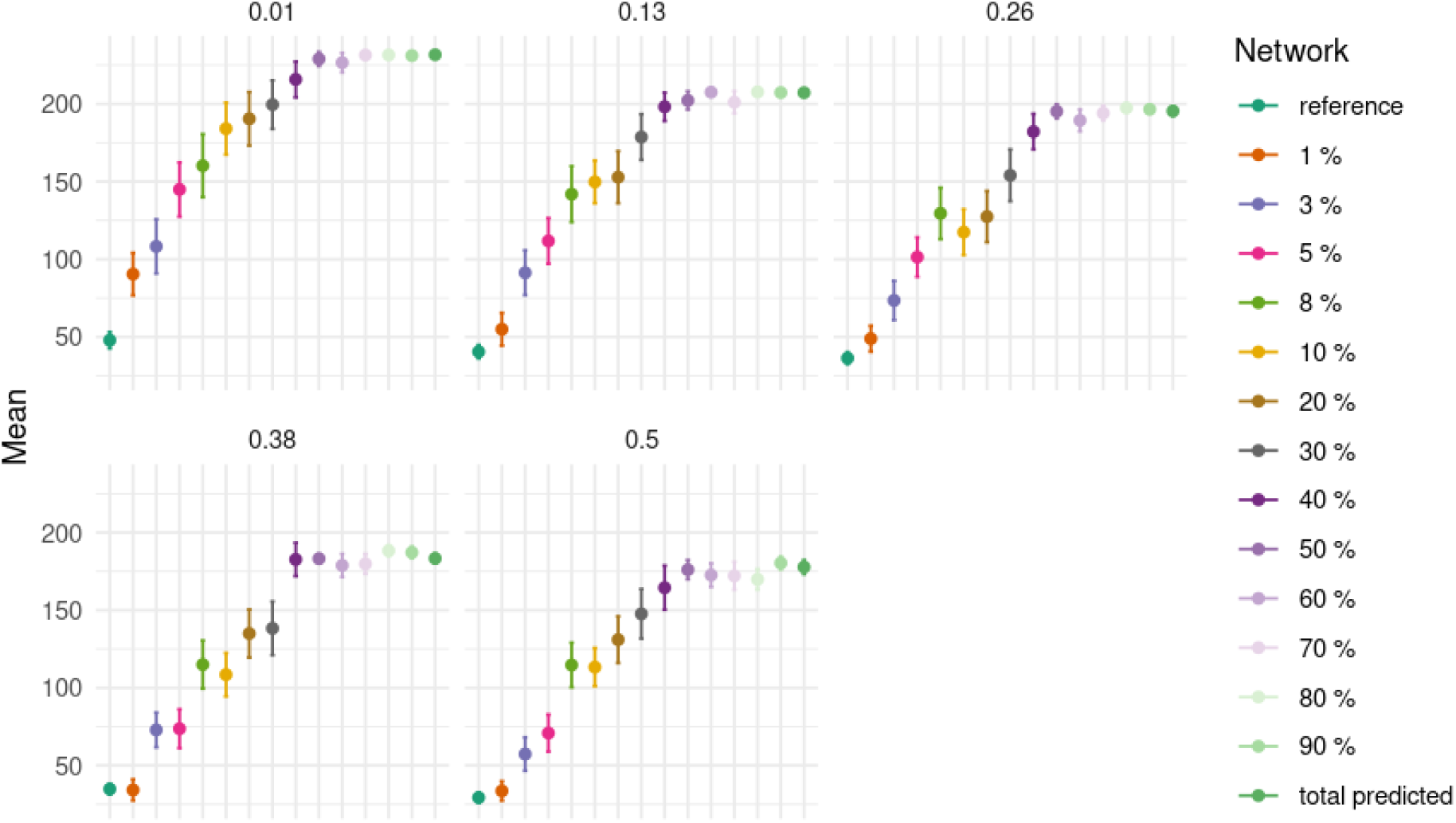
Incertitude study on the final epidemic size: *Pinf* fixed at 0.1 vs variation of *Prec*: case when the most probable link are added (*Top*_*d*_*ownscenario*)

In the *Bottom up scenario*, a similar pattern is observed when adding the least probable links (Figure 8). In other words, adding 1% of links with a low probability of existence does not impact the final size of the epidemic compared to the reference network. Still, this final size increases gradually as more links are added.

**Figure 8.**
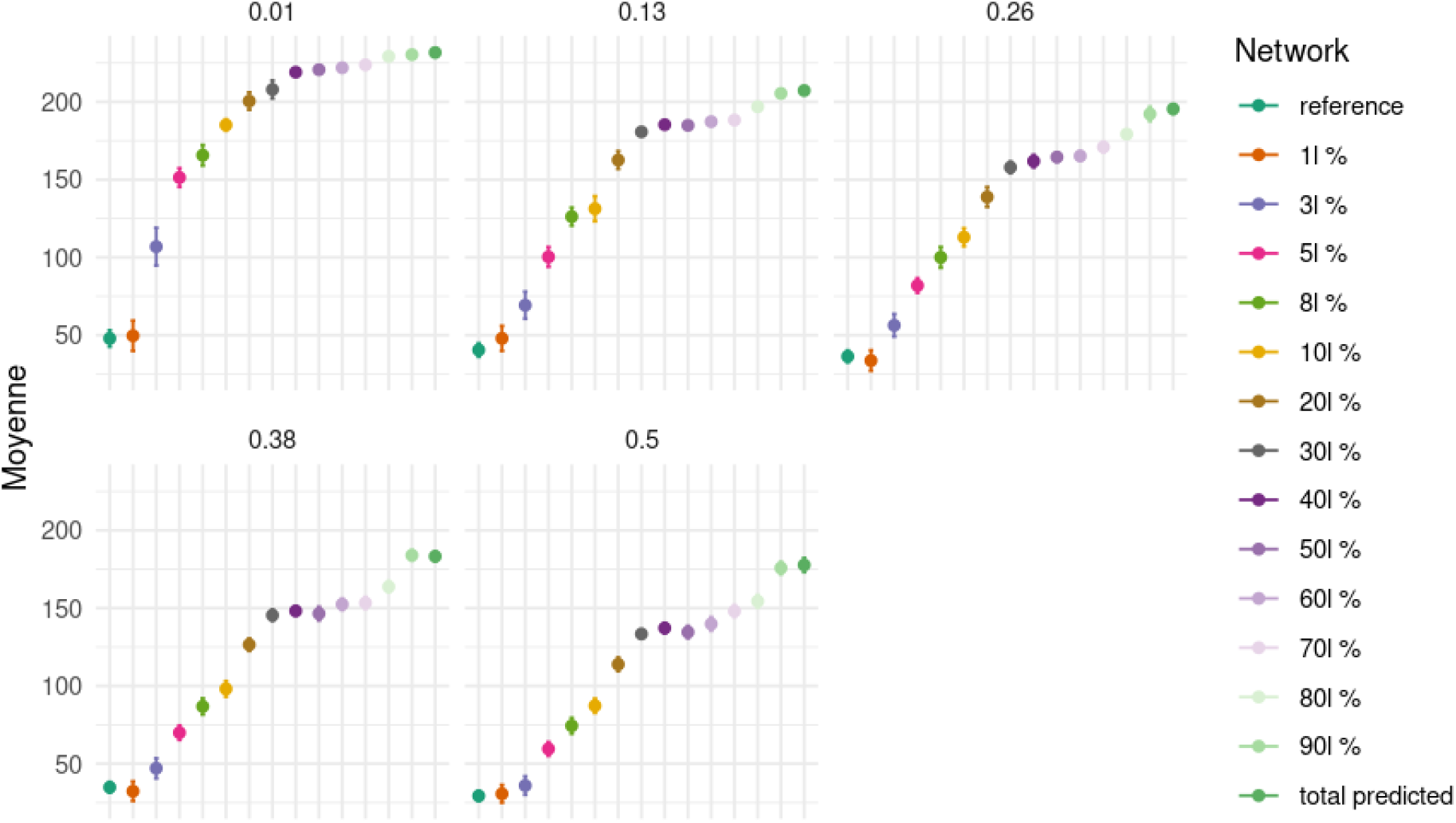
Incertitude study on the final epidemic size: *Pinf* fixed at 0.1 vs variation of *Prec*: case when the least probable link are added *Bottom*_*u*_ *pscenario*

Next, we analyzed starting from the fully predicted network. We observed that the absence of links between 10% and 50% does not impact the results significantly; the final size remains stable (Figure 7). However, removing less probable links changes the final size results; stability is maintained only when removing 10%, after which the results significantly differ (Figure 8).

Variation of the number and the identity of the vulnerable node, but the characteristics remain stable As mentioned above, the vulnerable nodes identified in the reference network vary according to the node groups. In the case where the most probable links are added, the number of identified vulnerable nodes generally follows the same pattern in most configurations (Figure 9): high in group Q1 (between 0 and 26), decreases in group Q2 (0 and 6), exceptionally 40 in one configuration, increases again in group Q3 (0 and 55), and reaches its maximum in group Q4 (17 and 59). The identity of the vulnerable nodes fluctuates.

**Figure 9.**
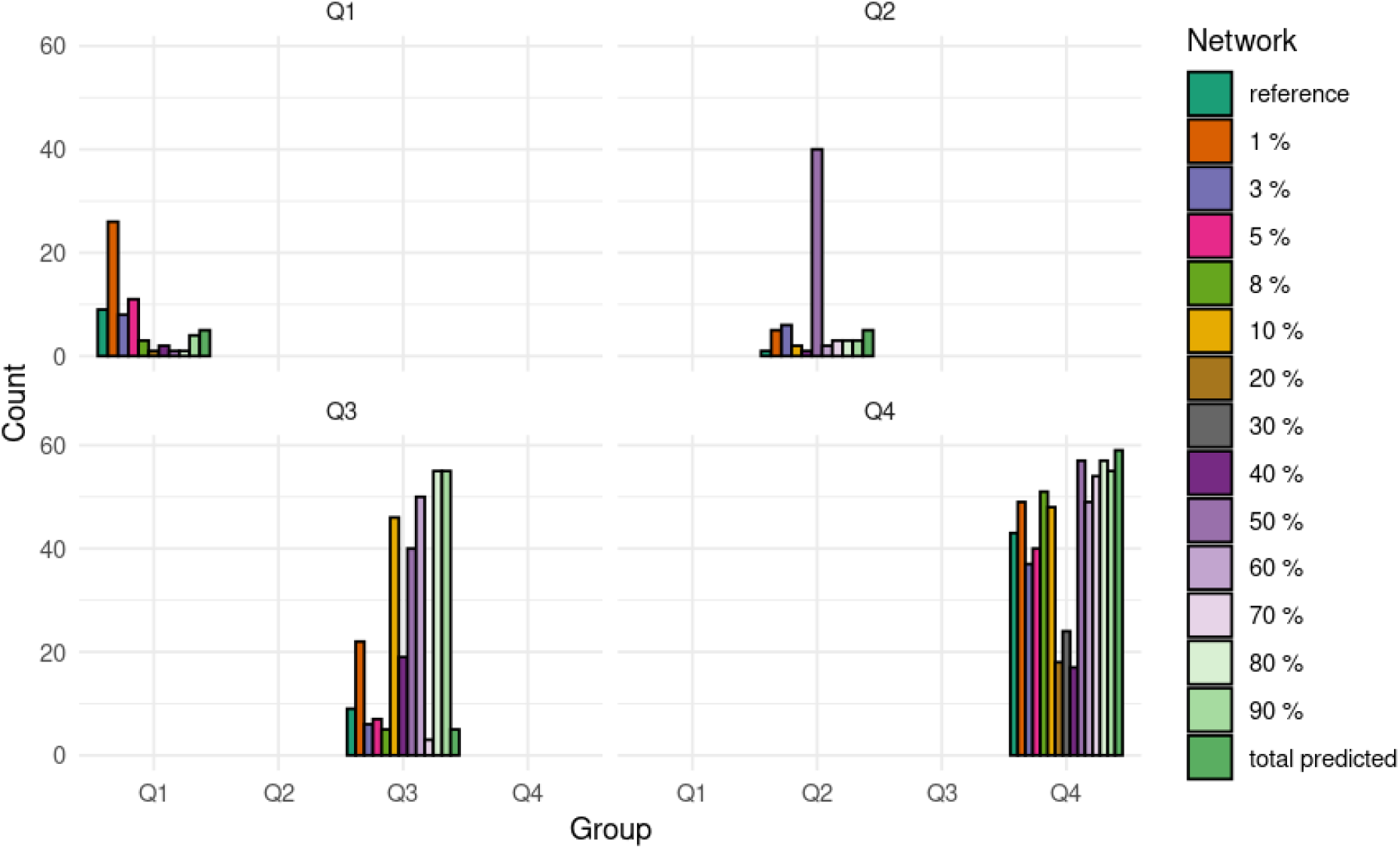
Number of high vulnerable node per network when ading graduaaly the most probable links

When adding the most probable links, and without considering transmission and recovery probabilities (groups combination), the number of vulnerable nodes per network varies between 19 (20%) and 138 (50%). The number of nodes in common between the reference network and the other networks varies between 8 and 38. The most vulnerable nodes in common between configuration 1% and 10% are between 21 and 38. Twenty nodes at 50% and less than 20 for the other configurations. The number of nodes in common between the total predicted network and the others varies between 6 and 57. There are 18 nodes in common between the total predicted network and the reference network. This number is raised from 50% (43), 60% (50), 70% (54), 80% (57), 90% (55).

Considering all the vulnerable nodes, 159 over 235 nodes will be vulnerable at least once in a given combination or network. Overall, for all possible combinations of Pinf and Prec, a maximum of 19 (network with 20% additional link) and 138 (network with 50% additional link) nodes are vulnerable.

The heatmap (Figure 10) reveals that in-closeness is consistently important across many configurations, followed by in-neighbourhood, which also frequently appears essential. In-degree, eigenvector, and in-h index are important but less often than in-closeness and in-neighbourhood. Conversely, in-strength rarely seems necessary, while rainfall, small ruminant population, and DMP (Dry Matter Product) are observed as significant even less frequently. Some structural characteristics, such as betweenness and transitivity, never appear as necessary. These results show that structural characteristics are more important than socio-economic characteristics, regardless of the networks.

**Figure 10.**
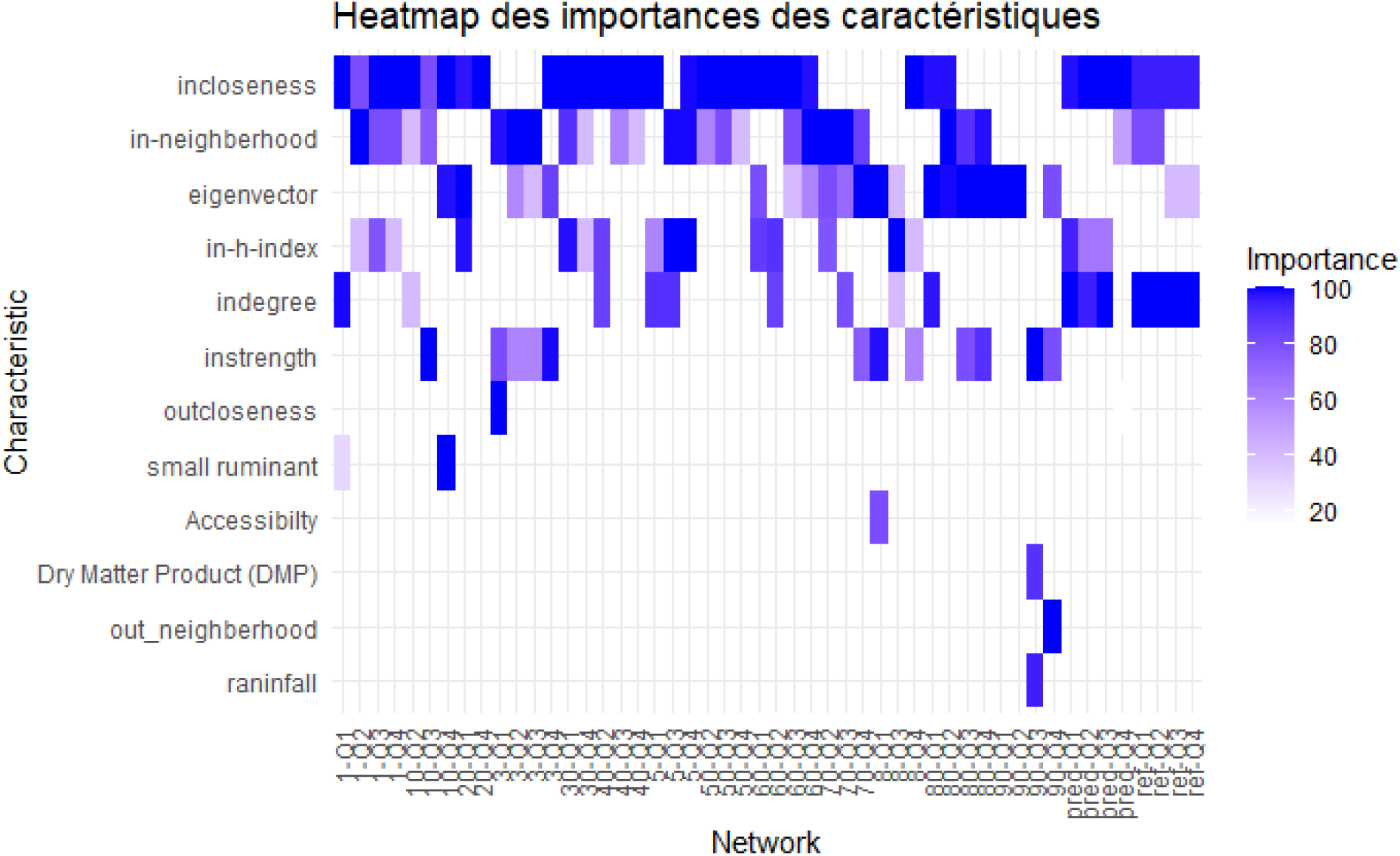
Heatmap of the most important characteristic of a vulnerable node in networks generated by adding most probable link

## Discussion

In this work, we relied on numerical simulations to analyse the pattern of PPR diffusion in Nigeria and identify characteristics of potential sentinel nodes. Additionally, it offers valuable insights into understanding PPR propagation and provides recommendations for improving surveillance and intervention strategies.

### Surveillance based on Contagion Clusters

We have identified group nodes with the same potential to infect each other. Interestingly, these groups are located in different states of Nigeria. This implies that if one node in such a group becomes infected, all other nodes in the same group (or community) will likely be rapidly infected, regardless of distance. Monitoring strategies focused on these contagion groups are essential to prevent rapid interstate disease spread. For example, the study by Salathé et al. (2010)^23^ demonstrated that in networks with strong community structures, vaccination interventions targeting individuals who connect to different communities are more effective than those targeting highly connected individuals. Based on these findings, we can hypothesize that targeted vaccination of nodes within these communities could be advantageous, particularly when vaccine supplies are limited.

Our study also revealed that eliminating the principal routes connecting these communities is insufficient to prevent the spread of PPR across the livestock mobility network. Restricting or prohibiting certain routes for herders or traders is unrealistic and could worsen the situation by encouraging alternative routes. An alternative approach could involve establishing random checkpoints along these routes. However, this measure remains impractical and burdensome for herders and traders. A more effective solution would be to position checkpoints at peripheral nodes. These nodes play an important role within their communities and serve as links to other communities, making them strategic points for monitoring and controlling disease spread.

### Targeted surveillance of PPR in Nigeria: The backbone serves as an ideal candidate

One of the most important results of this study is the role of the backbone and its nodes capability to serve as the best candidate in surveillance. The backbone is a weight- and connectivity-preserving subgraph that suffices to calculate all the shortest paths of weighted graphs^24^. Indeed, removing backbone nodes reduces key network performance metrics—such as the largest connected component (LCC), efficiency (Eff), and total flow (TF)—highlighting the backbone’s critical role in maintaining the network’s robustness. This aligns with the results of Correia (2023)^25^, which highlights the backbone as an important subgraph for the spread of epidemics, the robustness of social networks and any communication dynamics that depend on the shortest paths in a complex network. The backbone nodes are the network’s primary connectors, facilitating information flow. Their removal drastically reduces the LCC from 333 to 28 links, indicating a fragmentation of the network into isolated sub-networks. This fragmentation impedes and function of the network. The efficiency drop from 1.31*×*10^*−*3^ to 5.95 *×*10^*−*5^ after backbone removal illustrates how vital these nodes are to the network’s optimal performance. Backbone nodes, due to their high degree of connectivity, are likely to be infected early in an outbreak. Given the critical role that backbone nodes play in maintaining network connectivity and performance, they are ideal candidates for disease surveillance. Monitoring these nodes can indicate an epidemic’s presence and progression early. With fewer nodes to monitor than the entire network, focusing on backbone nodes allows for efficient allocation of resources and more intensive surveillance efforts. Health authorities can implement this by reallocating resources and health personnel or establishing health reporting at backbone nodes to track signs of disease outbreaks. Continuous data collection and analysis from these nodes can identify unusual patterns in disease that may indicate an outbreak, and establishing rapid response protocols can ensure swift containment and mitigation.

Furthermore, this study revealed, according to the classification made on the reference network, that in-closeness, in-degree, in-neighbourhood, and eigenvector centrality influence node vulnerability and are reliable according to the probability of transmission and recovery. According to Santiago Segarra et al. (2016)^26^, in-closeness, in-degree and eigenvector are stable measures on a weighted network. However, Lentz et al. (2016)^27^ noted that centrality measures are challenging to compute and can fluctuate significantly over time, depending on the specific period analyzed, as node rankings are not consistently stable^28^. Consequently, greater emphasis on rigorous data collection and careful timing of measurements is essential to ensure the reliability of these results.

### Contribution of link prediction to the reliability of results

This study also assessed the feasibility of using predicted links to complete a partial network. We investigated the uncertainty/reliability of the results, particularly about the final size of the epidemic. Adding 1% of links (approximately 46 links) to the reference network does not impact the propagation of PPR or the final epidemic size. This lack of impact can be attributed to the minimal increase in overall connectivity, insufficient to create new, significant pathways for disease spread. Essentially, adding a small percentage of links does not alter the network structure enough to affect the dynamics of epidemic propagation.

The network’s structural changes justify the observed variations in final epidemic size with the addition of links (between 3% and 40%). As Watts and Strogatz (1998)^29^ demonstrated, even small modifications (in our case, 3%) in network connectivity can significantly change dynamic processes such as epidemic spread. These changes further support the structural dependence of epidemic dynamics on network configuration. Adding links (up to 40%) increases the network’s connectivity, providing more pathways for disease spread, thus increasing the size of the epidemic. However, from 50% of the most probable links, the final size of the epidemic no longer varies significantly. However, this observation is not similar when the least probable links from the HRG model are added to the reference network. In fact, 90% more links are needed to reach a stable final epidemic size. A probable explanation lies in the prediction model that considers the hierarchical structure. When the model adds the most probable links, it selects connections that tend to form a hierarchical structure within the network. These links are chosen first because they help establish a well-organized, layered connection between nodes. However, when the model starts adding the least probable links, it focuses on those that are less likely to contribute to a hierarchical structure. These connections often have fewer connections and less influence, meaning they do not significantly affect the overall structure of the network. Essentially, the initial links are added to create a hierarchical framework, while the later, less probable links fill in gaps without altering the structure fundamentally. Therefore, epidemics spread almost instantaneously in networks with scale-free degree distributions^30^; this feature is associated with an epidemic propagation that follows precise hierarchical dynamics^30^. Once the highly connected hubs are reached, the infection pervades the network in a progressive cascade across smaller degree classes^30^. Another possible explanation is the concept of the percolation threshold^31^ in network theory can explain this plateau. Once the network connectivity is reduced below a critical threshold, the network becomes highly fragmented, preventing the epidemic from spreading effectively. The remaining network is no longer a single connected component but a collection of small, isolated sub-networks^32^. The networks exhibit a critical threshold of connectivity below which large-scale connectivity is lost. The percolation threshold is reached more quickly when most probable links are added, effectively increasing the network’s connectivity. In contrast, less probable links contribute weakly to the structure, delaying the point where large-scale propagation becomes possible. This could explain why stabilizing the epidemic size requires a much higher percentage of added links in the top down.

These results suggest that utilizing a partial network containing only 50% of the most probable predicted links can produce reliable outcomes. According to these results, we suggest adding 50% of the most probable links from the total predicted network to the partially observed network; this can improve the reliability of the results. In contrast, adding the least probable links takes up to 90% of the links for the epidemic size to stabilize. These low probable links are likely peripheral or redundant in the network. They might connect more isolated nodes or establish less important pathways, which initially don’t contribute much to the overall spread. As more of these links are added, they gradually enhance network connectivity but less efficiently, requiring more links (up to 90%) to achieve the same level of epidemic spread. These links create new pathways that eventually fill in gaps in the network, but the process is slower and less direct than adding the most probable links. This outcome also highlights that selecting links to complete a partial network should be done cautiously. It is essential to prioritize those with the highest probabilities, as they are more likely to contribute to a stable and predictable network structure and are more likely to exist. Adding lower-probable links can create unexpected routes and alter the system’s overall behaviour, potentially leading to less efficient or less stable network dynamics and links that probably don’t exist.

The variation in the number and identity of sentinel nodes across different parameter groups can be attributed to the dynamic nature of network properties and the stochasticity of our model. The network’s structure and the epidemic’s progression are highly sensitive to initial conditions and parameter variations, leading to different nodes being identified as highly vulnerable in each group.

However, specific structural characteristics, such as in-closeness and in-neighborhood, consistently demonstrate their importance across all parameter groups, highlighting their fundamental role in determining node vulnerability. Moreover, these characteristics remain robust even when the network structure changes, such as by adding links.

In contrast, while necessary, other centrality measures, such as in-degree, eigenvector centrality, and H-index, do not exhibit the same reliability or consistency across all generated network structures.

### Limitations and perspectives

Despite the robustness of our findings regarding the reliability of vulnerable node characteristics, this study has several limitations: Our analysis assumes a static network, whereas real-world networks are often dynamic, with links and nodes changing over time^33^, and backbone nodes can also evolve over time. However, methods that account for temporality in backbone extraction have been developed^34^,^35^ and can be used to improve the targeting of nodes. Furthermore, this static assumption limits the ability to capture temporal variations in network structure and interactions, which can influence epidemic dynamics.

The epidemic model used here simplifies the complexity of real-world disease transmission, which factors like animal population, species, age, environmental conditions and vaccination coverage. However, this simplification may not fully capture the multifaceted nature of disease spread. While we analyzed nodes across a few states, the study is still limited to a specific region, potentially reducing the generalizability of our findings to other geographical contexts. Networks in other areas may exhibit different structural and epidemiological characteristics.

In our future work, we will extend the analysis to dynamic networks and use more complex models that account for additional factors such as vaccination rate, seasonality, and animal population, which could yield more accurate predictions. We will expand the geographical scope to include diverse regions. This broader analysis can reveal how different environmental, seasonality, socioeconomic, and cultural factors influence network structure, epidemic dynamics, and node vulnerability.

## Methods

### Reference Network and identification of contagion cluster

#### Reference Network representation

Based on a preliminary analysis conducted by partners of the Lidiski project, a total of ten markets in three Nigerian states were selected for sampling. 6 markets were sampled in Plateau, and for security, logistic and accessibility reasons, only two markets were sampled in both of Bauchi and Kano States (see details in^20^). Data on origin and destination and the number of animals moved is collected. From this data, we reconstructed the mobility network (reference network or observed network) whose nodes are villages, and links represnt movemnts between two villages weighted the number of heads exchanged. The generated weighted network has 233 nodes and 335 links. Among these movements, 184 have villages as origins and destinations in the 3 states of the study area. This network is considered partial because data collection was conducted only once, with a limited number of farmers (100 respondents per market) and in a limited number of markets, making it non-exhaustive.

#### Identification of contagion cluster

This step follows COCLEA algorithm (COntagion CLusters Extraction Algorithm)^16^)to identify infection clusters, which correspond to a group of nodes that can mutually infect each other by leveraging community detection techniques. This approach is differnt from previous communities based algoithms that were based on modularity optimisation, since its advantage lies in its use of dynamics, specifically the SIR model and network reliability.COCLEA identifies these “contagion clusters” by ensuring that each cluster induces a strongly connected component (SCC) in the network and that connectivity within the cluster is strong enough for significant mutual influence. The identification of these clusters could help in devising target interventions more effectively, potentially limiting the spread of infection. By understanding how infections propagate within and between communities, we can develop more informed strategies for disease control and prevention.

Following the approach of Nath (2019)^16^, we varied the cluster size n, which corresponds to the desired number of nodes in the strongly connected component (SCC), using values n=2,5,10,20,30, and V, that correspond to the total number of network (n = 233). We then compared the number of detected communities and their distribution areas. Based on this comparison, the optimal cluster size n was selected by choosing the best balance, determined as the most appropriate compromise between these n.

### Simulation of PPR-like disease in the Reference network

#### PPR spread using the SIR model

We simulated the spread of PPR disease through animal movements between the villages of the three Nigerian States using a stochastic Susceptible-Infectious-Recovered (SIR) weighted model:

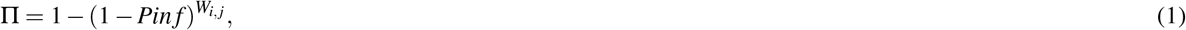

*Pin f* represents the probability that an animal from an infected village is infectious and transmits the disease to other animals in susceptible areas, and *W*_*i*, *j*_, represents the number of animals moved from an infected node *j* to *i*. Intra-village transmission is neglected in this study, as PPR is a highly contagious infectious disease that spreads rapidly through animal movements^14^. Initially, all nodes are susceptible, except for one village with a non-zero number of outgoing links. At each simulation stage, an infected village, *i*, can transmit the disease to all its neighbors in a susceptible state. Once infected, these neighbours can spread the disease to other villages. An infected village reaches the state R with the probability *Prec*. This status represents a village recovered from PPR and cannot be reinfected because this disease confers lifelong immunity.

Choosing transmission *Pinf* and recovery probabilities *Prec* parameters is crucial yet often non-trivial during disease simulations. While estimates exist for these parameters at an individual level, their variability on a geographical scale is not always clearly defined. These parameter values can vary significantly depending on numerous factors such as demographic characteristics, individual behaviors, and local environmental conditions. Additionally, the transmission and recovery rates for peste des petits ruminants (PPR) can vary due to various factors including viral strain, environmental conditions, and characteristics of infected animals. Testing an extensive range of values allows us to explore the sensitivity of our models to variations in these parameters. Therefore, we explored 10,000 combinations of *Pinf* (between 0.00001 and 0.2) and *Prec* (between 0.01 and 0.5). Following exploratory analyses regarding the average final epidemic size variability, we conducted 300 simulations for each combination of the two parameters, resulting in a total of 10,000*300 simulations. Simulations were run on a weekly time step for 50 weeks. We conducted a simple sensitivity analysis using contour plots to identify groups of parameter combinations that behave similarly regarding the final size of the epidemic, then we applied quartiles to facilitate the comparison between the parameter groups.

#### Identification of sentinel nodes

Nodes frequently infected before the epidemic’s peak are deemed vulnerable and can serve as sentinel nodes. Initially, the optimal distribution of the infection frequency of nodes before the peak is verified. The preferred model is found to be the lognormal distribution. After fitting the lognormal distribution, estimated parameters—mean (mu) and standard deviation (sigma)—are derived. The threshold selection relies on a quantile of the lognormal distribution, particularly the 75th percentile. Based on this threshold, three statuses were defined: if a node is affected before the epidemic peak with a frequency higher than the threshold, it is considered highly vulnerable. If a node is frequently affected more before than after the peak, but with a frequency below the threshold, it is classified as lowly vulnerable. Lastly, if a node is frequently affected after the epidemic peak, it is considered infected but not vulnerable. Here, nodes identified as highly vulnerable are considered the best candidates to be sentinel node.

#### Structural, socio-economics and environmental characteristics of sentinel node

The estimation conducted in the previous step classifies nodes into three categories: high vulnerable, low vulnerable, infected but not vulnerable (Infec-NV). Centrality measures (in/out degree, in/out closeness, betweenness, eigenvector, in/out neighborhood, in/out H index, transitivity, in/out strength) were estimated for each node. Socio-economic and environmental variables were extracted for each node. Human population data for 2020 were obtained in raster format from https://data.humdata.org/dataset/worldpop-population-counts-for-nigeria. The sum and average per ward were then extracted. Additionally, goat and sheep populations for 2015 were acquired in raster form from https://dataverse.harvard.edu/dataverse/glw4. The sum and average per ward and species were extracted, and the populations were subsequently merged into small ruminants. In Copernicus, Rainfall and DMP (Dry Matter Product) from https://land.copernicus.eu/global/products/dmp, representing the overall growth rate or increase in vegetation dry biomass, was utilized. Raster data from 1999 to 2020 were extracted, and the median for this period was estimated. Subsequently, an extraction by ward was performed. Moreover, accessibility or travel time (in minutes) to the nearest city in 2015 was obtained from https://data.malariaatlas.org/. Centrality measures and socio-economic and environmental variables were employed as features in the random forest classification method to identify the variables that explain vulnerability.

### Impact of target node (identified as sentinel) or bridge or backbone removal on PPR propagation

In this section, we explored various scenarios that could benefit from improving the surveillance and control of PPR. Previously, we identified vulnerable nodes that could be good candidates for sentinel nodes. We simulated surveillance by removing these nodes. The underlying hypothesis is that if surveillance and control are focused on these nodes, disease spread will be limited from these critical points. Since vulnerable nodes have been identified for each parameter group, this removal involves all sets of nodes identified as vulnerable within each group. Additionally, we identified communities or contagion clusters connected by “bridge” links. These “bridge” links refer to the critical connections between the communities within a network. We considered that surveillance could be effective by focusing on these “bridges” and thus tested removing these links.

Another scenario tested is the removal of backbone nodes in the network. In this study^18^, 45% of the backbone nodes appear vulnerable, suggesting that the backbone could be a good candidate for surveillance. After the extraction of backbone using the disparity filter^19^, the impact of these removals on the network was tested, we used measures such as LCC (Largest Connected Component), Eff (Efficiency) and TF (Total Flow)^21^ These measures allowed us to evaluate the impact of these removals on the network’s robustness. We also used the final epidemic size to measure the removal’s impact on the spread of PPR.

### Uncertainty analysis

#### Prediction of missing links

The reconstructed network represents only a fraction of the potential movements that should be present. Analyzing this incomplete data can significantly impact estimates of network-level statistics and inferences about the network’s structural properties. To enhance the prediction of the animal mobility network reconstruction, first, several methods were tested considering edge weight for methods:

i. Neighbourhood-based predictors^36^: which assign high likelihood scores to node pairs that share many neighbours
ii. CAR-based predictors^37^: which assign high likelihood scores to node pairs that share many neighbours that also interact with themselves
iii. The Hierarchical Random Graph method (HRG)^38^ searches the space of all possible dendrograms of a network for the ones that best fit its hierarchical structure. Non-adjacent node pairs that have high average probability of being connected within these dendrograms are considered good candidates for interaction
iv. The Structural Perturbation Method (SPM)^39^ is based on the hypothesis that links are predictable if removing them has only small effects on network structure.

The performance of these link predictors was assessed by pruning edges from the network and checking whether the methods give high likelihood scores to the removed edges. Performance is assessed using the AUC metric. All these sections were done using the R package “LinkPrediction.” The best method (with AUC = 0.9) was the HRG model. Then, socio-economic and environmental variables were employed from these predicted links to predict the weight between each expected link. As in Garcia Garcia et al.^40^, a hurdle model, the negative binomial distribution was applied because of a significant over-dispersion. We predicted the weights of our links, and the best predictive model considers the following variables (according to the lowest Akaike Information Criterion(AIC)): distance between origin and destination (estimated using the latitude and longitude), small ruminant population at origin and destination, accessibility at origin and destination, and dry matter product at origin.

#### Simulation of PPR in predicted network

Simulations of PPR were conducted on 14 representative networks using specific parameter values of *Pinf* and *Prec* from four predefined groups. Links were added to the reference network based on their probability of existence. Here, we have carried out two situations; the first, we have added the links that have a greater possibility of existing, i.e. with a higher probability. In the 2nd situation, on the other hand, we have added the links that are less likely to exist, i.e. those with a low probability. Initially, 1% of the most probable links were added, followed by 3%,5%, 8%, 10%, 20%, 30%, 40%, 50%, 60%, 70%, 80%, 90%, up to the fully predicted network 100%. In this way, 14 additional networks were constructed by adding the least probable links. The average final size of the epidemics was estimated, and the number and identity of vulnerable nodes over all simulations was then compared with the reference network.

After conducting the simulations, we followed the same procedures outlined earlier, specifically focusing on identifying vulnerable nodes and their characteristics. The objective is to determine the effects of adding different types of links to the reference network and to observe any changes in the results.

LaTeX formats citations and references automatically using the bibliography records in your .bib file, which you can edit via the project menu. Use the cite command for an inline citation, e.g.^**?**^. For data citations of datasets uploaded to e.g. *figshare*, please use the `howpublished` option in the bib entry to specify the platform and the link, as in the `Hao:gidmaps:2014` example in the sample bibliography file.

## Acknowledgements

The research was funded by a grant from the European Commission (Development Cooperation Instruments) awarded to the project ‘EU Support to Livestock Disease Surveillance Knowledge Integration -LIDISKI’(FOOD/2019/410-957).

## Author contributions statement

Conceived and designed the analysis: AA, MA, AM, SE; Collected and curated the data: SI, MB; Performed the analysis: AM; Wrote the draft paper: AM; Critical review: AA, MA, MC, EC.

## Additional information

**Table 2.**
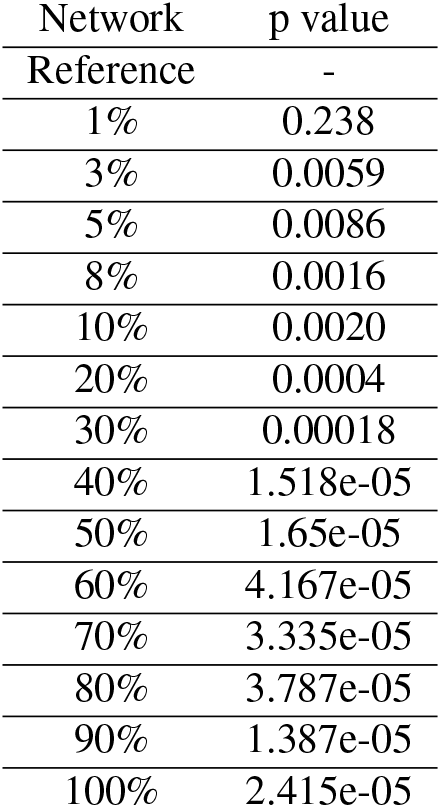
Difference Between Epidemic Size and Final Size with Networks Generated from the Addition of the most probable Links predicted by the HRG model.

## References

1. Clemmons, E. A., Alfson, K. J. & Dutton, J. W. Transboundary animal diseases, an overview of 17 diseases with potential for global spread and serious consequences. 11, 2039, DOI: 10.3390/ani11072039. Number: 7 Publisher: Multidisciplinary Digital Publishing Institute.

2. Progress on PPR control and eradication strategy.

3. Njeumi, F., Bailey, D., Soula, J. J., Diop, B. & Tekola, B. G. Eradicating the scourge of peste des petits ruminants from the world. 12, 313, DOI: 10.3390/v12030313. Number: 3 Publisher: Multidisciplinary Digital Publishing Institute.

4. Kabir, A. Peste des petits ruminants: A review. 8, DOI: 10.19045/bspab.2019.80063.

5. Torsson, E. et al. History and current status of peste des petits ruminants virus in tanzania. 6, 32701, DOI: 10.3402/iee.v6.32701. Publisher: Taylor & Francis _eprint: https://doi.org/10.3402/iee.v6.32701.

6. Idoga, E. S. et al. A review of the current status of peste des petits ruminants epidemiology in small ruminants in tanzania. 7, DOI: 10.3389/fvets.2020.592662. Publisher: Frontiers.

7. Baron, M. D., Diallo, A., Lancelot, R. & Libeau, G. Chapter one - peste des petits ruminants virus. In Kielian, M., Maramorosch, K. & Mettenleiter, T. C. (eds.) Advances in Virus Research, vol. 95, 1–42, DOI: 10.1016/bs.aivir.2016.02.001 (Academic Press).

8. Diallo, A. La peste des petits ruminants : une maladie longtemps ignorée. DOI: 10.4267/2042/47951. Publisher: Persée - Portail des revues scientifiques en SHS.

9. Baazizi, R. et al. Peste des petits ruminants (PPR): A neglected tropical disease in maghreb region of north africa and its threat to europe. 12, e0175461, DOI: 10.1371/journal.pone.0175461. Publisher: Public Library of Science.

10. Apolloni, A. et al. Towards the description of livestock mobility in sahelian africa: Some results from a survey in mauritania. 13, e0191565, DOI: 10.1371/journal.pone.0191565. Publisher: Public Library of Science.

11. Apolloni, A., Corniaux, C., Coste, C., Lancelot, R. & Touré, I. Livestock mobility in west africa and sahel and transboundary animal diseases. In Kardjadj, M., Diallo, A. & Lancelot, R. (eds.) Transboundary Animal Diseases in Sahelian Africa and Connected Regions, 31–52, DOI: 10.1007/978-3-030-25385-1_3 (Springer International Publishing).

12. Nicolas, G. et al. Predictive gravity models of livestock mobility in mauritania: The effects of supply, demand and cultural factors. 13, e0199547, DOI: 10.1371/journal.pone.0199547. Publisher: Public Library of Science.

13. Colizza, V. & Vespignani, A. Epidemic modeling in metapopulation systems with heterogeneous coupling pattern: theory and simulations. 251, 450–467, DOI: 10.1016/j.jtbi.2007.11.028.

14. Bajardi, P., Barrat, A., Savini, L. & Colizza, V. Optimizing surveillance for livestock disease spreading through animal movements. 9, 2814–2825, DOI: 10.1098/rsif.2012.0289.

15. Chaters, G. et al. Analysing livestock network data for infectious disease control: An argument for routine data collection in emerging economies. 374, 20180264, DOI: 10.1098/rstb.2018.0264.

16. Nath, M. et al. Using network reliability to understand international food trade dynamics. 812, 524–535.

17. Mishra, R., Eubank, S., Nath, M., Amundsen, M. & Adiga, A. Community detection using moore-shannon network reliability: Application to food networks. In Cherifi, H., Mantegna, R. N., Rocha, L. M., Cherifi, C. & Micciche, S. (eds.) Complex Networks and Their Applications XI, Studies in Computational Intelligence, 271–282, DOI: 10.1007/978-3-031-21131-7_21 (Springer International Publishing).

18. Mesdour, A. et al. Assessing the impact of structural modifications in the construction of surveillance network for transboundary animal diseases: the role of backbone and sentinel nodes. DOI: 10.1101/2024.04.24.590906.

19. Neal, Z. P. backbone: An r package to extract network backbones. 17, e0269137, DOI: 10.1371/journal.pone.0269137.2203.11055[cs].

20. Ijoma, S. I. et al. Combining market surveys and participatory approaches to map small ruminant mobility in three selected states in northern nigeria, DOI: 10.21203/rs.3.rs-5122130/v1. ISSN: 2693-5015.

21. Bellingeri, M. et al. Link and node removal in real social networks: A review. 8.

22. Latora, V. & Marchiori, M. A measure of centrality based on network efficiency. 9, 188, DOI: 10.1088/1367-2630/9/6/188.

23. Salathé, M. & Jones, J. H. Dynamics and control of diseases in networks with community structure. 6, e1000736, DOI: 10.1371/journal.pcbi.1000736. Publisher: Public Library of Science.

24. Serrano, M., Boguñá, M. & Vespignani, A. Extracting the multiscale backbone of complex weighted networks. 106, 6483–6488, DOI: 10.1073/pnas.0808904106. Publisher: Proceedings of the National Academy of Sciences.

25. Correia, R. B., Barrat, A. & Rocha, L. M. Contact networks have small metric backbones that maintain community structure and are primary transmission subgraphs. 19, e1010854, DOI: 10.1371/journal.pcbi.1010854. Publisher: Public Library of Science.

26. Segarra, S. & Ribeiro, A. Stability and continuity of centrality measures in weighted graphs. 64, 543–555, DOI: 10.1109/TSP.2015.2486740. Conference Name: IEEE Transactions on Signal Processing.

27. Lentz, H. H. K. et al. Disease spread through animal movements: A static and temporal network analysis of pig trade in germany. 11, e0155196, DOI: 10.1371/journal.pone.0155196. Publisher: Public Library of Science.

28. Konschake, M., Lentz, H. H. K., Conraths, F. J., Hövel, P. & Selhorst, T. On the robustness of in- and out-components in a temporal network. 8, e55223, DOI: 10.1371/journal.pone.0055223. Publisher: Public Library of Science.

29. Watts, D. J. & Strogatz, S. H. Collective dynamics of ‘small-world’ networks. 393, 440–442, DOI: 10.1038/30918. Publisher: Nature Publishing Group.

30. Barthélemy, M., Barrat, A., Pastor-Satorras, R. & Vespignani, A. Velocity and hierarchical spread of epidemic outbreaks in scale-free networks. 92, 178701, DOI: 10.1103/PhysRevLett.92.178701. Publisher: American Physical Society.

31. Mohseni-Kabir, A., Pant, M., Towsley, D., Guha, S. & Swami, A. Percolation thresholds for robust network connectivity. J. Stat. Mech. Theory Exp. 2021, 013212, DOI: 10.1088/1742-5468/abd312 (2021).

32. Newman, M. E. J. Percolation and network resilience: A discussion of one of the simplest of processes taking place on networks, percolation, and its use as a model of network resilience. In Newman, M. (ed.) Networks: An Introduction, 0, DOI: 10.1093/acprof:oso/9780199206650.003.0016 (Oxford University Press).

33. Holme, P. & Saramäki, J. Temporal networks. 519, 97–125, DOI: 10.1016/j.physrep.2012.03.001.

34. Nadini, M., Bongiorno, C., Rizzo, A. & Porfiri, M. Detecting network backbones against time variations in node properties. 99, 855–878, DOI: 10.1007/s11071-019-05134-y.

35. Surano, F. V., Bongiorno, C., Zino, L., Porfiri, M. & Rizzo, A. Backbone reconstruction in temporal networks from epidemic data. 100, 042306, DOI: 10.1103/PhysRevE.100.042306.

36. Kerrache, S., Alharbi, R. & Benhidour, H. A scalable similarity-popularity link prediction method. 10, 6394, DOI: 10.1038/s41598-020-62636-1. Publisher: Nature Publishing Group.

37. Cannistraci, C. V., Alanis-Lobato, G. & Ravasi, T. From link-prediction in brain connectomes and protein interactomes to the local-community-paradigm in complex networks. 3, 1613, DOI: 10.1038/srep01613. Publisher: Nature Publishing Group.

38. Clauset, A., Moore, C. & Newman, M. E. J. Hierarchical structure and the prediction of missing links in networks. 453, 98–101, DOI: 10.1038/nature06830. Publisher: Nature Publishing Group.

39. Toward link predictability of complex networks | PNAS.

40. García García, K. M. et al. Environmental and economic determinants of temporal dynamics of the ruminant movement network of senegal. 13, 14482, DOI: 10.1038/s41598-023-40715-3. Number: 1 Publisher: Nature Publishing Group.

